# Biological meaning in protein embedding space is resolution-dependent

**DOI:** 10.64898/2026.06.12.730559

**Authors:** Licheng Zong, Jinzheng Ren, Yu Li, Robert D. Finn, Jiawei Wang

**Author notes:** These authors contributed equally to this work (Licheng Zong, Jinzheng Ren). Correspondence (J.W.), (R.D.F) and (Y.L.).

## Abstract

Protein language model embeddings are increasingly used to organise biological sequences, yet how biological meaning is encoded within embedding neighbourhoods remains poorly understood. Using two independent hierarchical enzyme systems, carbohydrate-active enzymes and peptidases, we investigated how biological interpretation changes across embedding organisations aligned to different levels of biological hierarchy. Different embedding organisations give rise to distinct neighbourhood semantics. When aligned to membership-boundary resolution, embeddings robustly separated artefacts and unrelated proteins from members of the target category. However, embeddings aligned to functional-grouping resolution maintained compositional neighbourhood structure for multi-domain proteins spanning more than one functional or catalytic group. Finally, embeddings aligned to local-family resolution recovered compact family-like neighbourhoods, including families withheld from training, while weakening broader membership-boundary and functional-grouping relationships. Moreover, embeddings optimised toward the same level of biological organisation retain different biological relationships depending on optimisation trajectory employed. Together, our results show that proximity in protein embedding space has no fixed biological interpretation. Instead, biological meaning emerges across embedding resolutions through selective preservation of different forms of biological organisation.

## Introduction

Protein language models (PLMs) have transformed biological sequence analysis by learning general-purpose embeddings from large protein sequence collections, capturing signals related to structure, function, evolution and biochemical properties^1–4^. These embeddings now underpin applications ranging from structure prediction and functional annotation to protein engineering and generative design^3, 5–9^. As protein embeddings become increasingly used to explore biological sequence space, local neighbourhoods have become a practical basis for inferring relationships between proteins. This raises a conceptual question: what biological relationship does the neighbourhood proximity represent?

Protein embeddings are commonly developed and assessed in the context of functional prediction, retrieval and representation quality. Task-adapted and multimodal models shape embeddings to improve prediction or retrieval, while representation-level studies query whether embeddings are biologically informative or reliable across models and tasks^6, 8–11^. These approaches establish the value of protein embeddings for prediction, retrieval and representation-level assessment, but a local neighbourhood in the embedding space is a relational object, namely its biological meaning depends not only on what information a representation contains, but also on which relationships between proteins are preserved by the geometry of the space. This distinction reflects the layered and partially overlapping organisation of protein biology. Proteins can be related through local motifs, domains, structural folds or broader biological systems, such as functional classes^12–14^. These relationships often correlate, but they do not necessarily map one-to-one. Consequently, local proximity in an embedding space should be interpreted in relation to the biological information that the model has been trained, adapted or otherwise shaped to preserve. Nearby proteins may share family-level, structural, functional or domain similarity, but these interpretations are not interchangeable. Protein embedding neighbourhoods therefore require explicit biological framing: proximity should be interpreted with respect to the relationship preserved by the embedding geometry and the biological level at which that relationship is defined. We therefore introduce protein embedding resolution as an organising principle for interpreting biological neighbourhoods in embedding space (Fig. 1). We define embedding resolution as the biological scale at which neighbourhood relationships can be meaningfully interpreted. Under this paradigm, embedding spaces are not universal maps of biological similarity, but organised geometries that selectively preserve some biological relationships while compressing or obscuring others. The same local proximity can therefore support different biological interpretations depending on the resolution at which the space is organised.

**Fig. 1.**
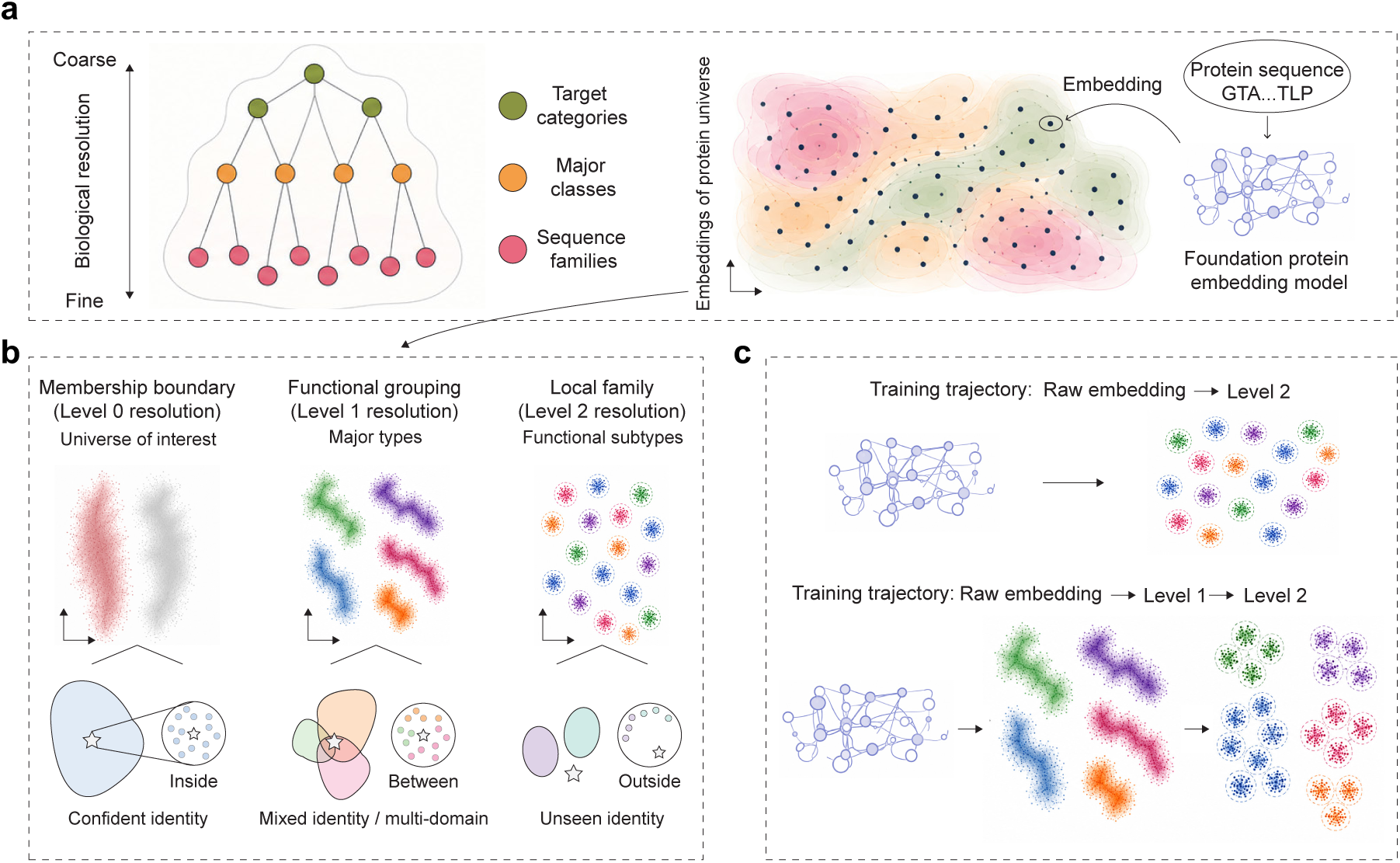
A resolution and trajectory framework for interpreting protein embedding neighbourhoods. **(a)** Schematic overview of protein embedding resolution. Biological organisation can be considered across multiple scales, from broad target category to intermediate major classes and fine-grained sequence families (left panel). Foundation protein language models encode protein sequences into continuous embedding spaces in which signals from different biological scales may coexist (right panel). **(b)** Resolution-matched embedding spaces used in this study. Level 0 represents broad membership boundary and is used to evaluate whether proteins lie inside or outside a protein category of interest. Level 1 represents intermediate functional categories and is used to examine relationships between major functional types, including proteins associated with more than one category. Level 2 represents fine-grained sequence-family organisation and is used to examine local-family level neighbourhoods, including families withheld from training. **(c)** Schematic of trajectory-dependent embedding organisation. Embeddings can be aligned directly from the pretrained representation to a family-level space, or sequentially through an intermediate Level 1 space before Level 2 alignment. These alternative paths provide a way to test whether the geometry reached at the same nominal resolution depends on the training trajectory.

To demonstrate this, we investigated this paradigm using two independent hierarchical enzyme systems: carbohydrate-active enzymes (CAZymes) and peptidases. CAZymes mediate carbohydrate recognition, modification and degradation, whereas peptidases mediate proteolysis^15–17^, essential for functions such as protein turnover and pathogenesis. Together, these enzyme categories are functionally diverse, data-rich and well annotated, and can account for substantial fractions of protein-coding genes in genomes (up to ∼5% for each category). CAZy and MEROPS provide curated label hierarchies in which the labels themselves carry explicit biological semantics: fine-grained sequence families are organised into broader, superfamily-like functional or catalytic groupings above the family level. We further introduced an outer membership boundary level by contrasting proteins inside and outside each target enzyme category. These semantically defined hierarchies allowed us to align the same pretrained foundation-model representation to three semantically matched biological resolutions: membership-boundary resolution, which determines whether a protein lies inside or outside the target category; functional-grouping resolution, which evaluates the broad functional or catalytic group a neighbourhood reflects; and local-family resolution, which evaluates whether specific family-like neighbourhood is preserved. We then used the resulting protein embedding spaces to test which biological relationships are preserved, lost or transformed at each embedding resolution, whether these patterns generalise across these two independent biological systems, and whether the optimisation trajectory used to reach a given resolution influences the resulting geometry. Together, this design allowed us to examine protein embedding space as resolution-dependent: neighbourhood meaning is not assumed from proximity alone, but evaluated through the biological relationships retained or reorganised at each resolution.

## Results

### Resolution-matched supervision reorganises CAZyme embedding space across biological scales

We tested the hypothesis that biological meaning in protein embedding space is resolution-dependent by organising embeddings around three different biological scales: membership boundary, functional grouping and specific enzyme family (Fig. 1a,b). We first applied this framework to carbohydrate-active enzymes (CAZymes), using the curated CAZy hierarchy as a semantically defined label system (Fig. 2a, b). CAZy provides fine-grained sequence-family annotations and groups these families into broader carbohydrate-active classes, including Glycoside Hydrolases (GH), Glycosyl Transferases (GT), Polysaccharide Lyases (PL), Carbohydrate Esterases (CE), Auxiliary Activities (AA, and a more general class) and Carbohydrate-Binding Modules (CBM), which span distinct functions associated with carbohydrate metabolism^15, 16^ (Fig. 2b). We extended this hierarchy with an outer membership-boundary set from Swiss-Prot^18^ to distinguish CAZymes from non-CAZyme proteins. This produced three biologically interpretable reference resolutions: membership-boundary resolution, corresponding to CAZyme versus non-CAZyme discrimination; functional-grouping resolution, corresponding to six major CAZy class organisation; and local-family resolution, corresponding to fine-grained CAZy family organisation.

**Fig. 2.**
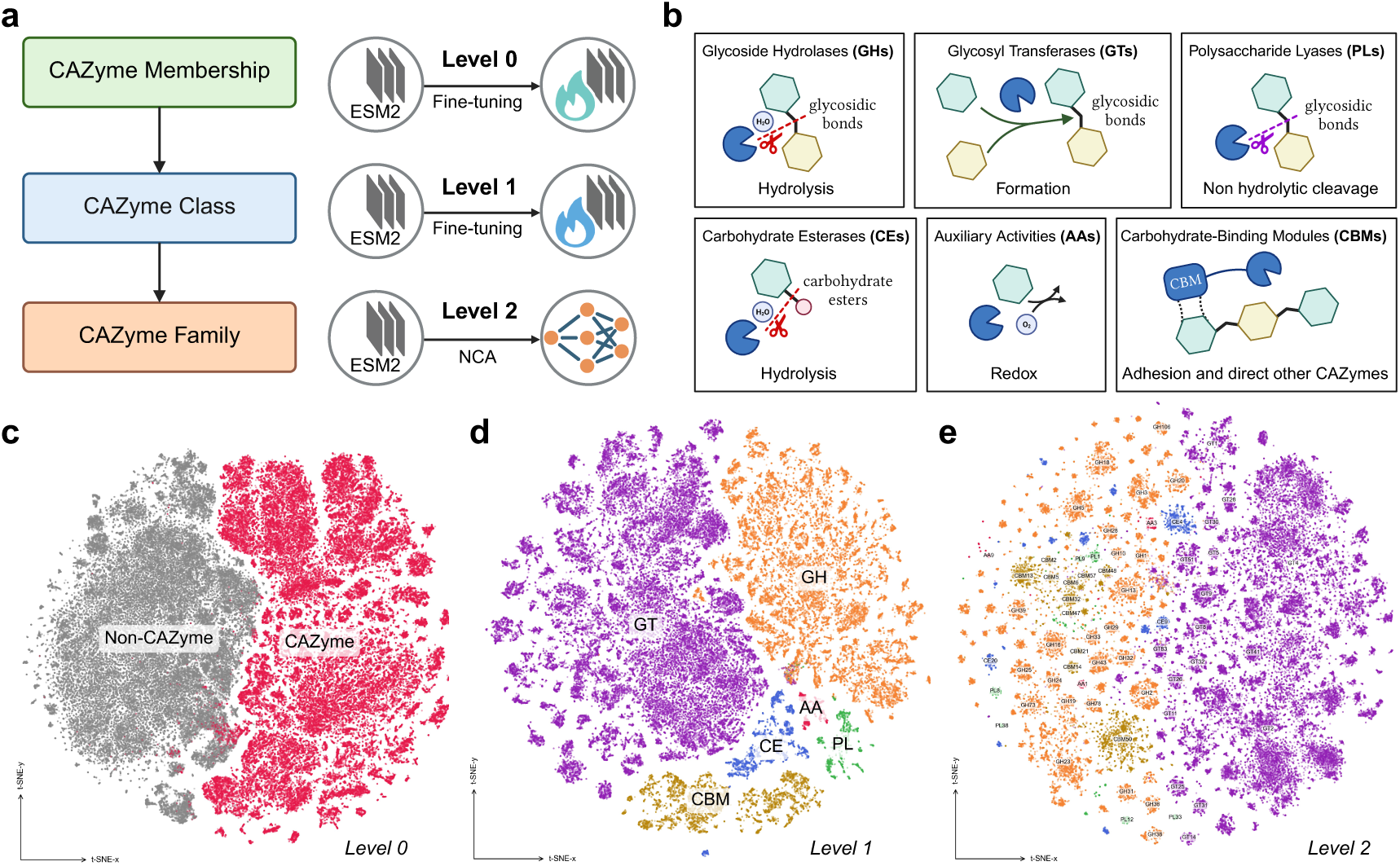
CAZymes as a three-level supervision testbed for resolution-matched embedding. **(a)** Carbohydrate-active enzymes (CAZymes) are organised at three biological resolutions used as supervision targets: CAZyme membership (Level 0, binary CAZy versus non-CAZy), CAZy class (Level 1, six classes), and CAZy family (Level 2, 416 families). Starting from pretrained ESM-2 representations, embeddings were aligned to each resolution through level-matched training: binary fine-tuning at L0, six-class fine-tuning using single-class CAZymes at L1, and Neighbourhood Components Analysis (NCA) family-level metric learning at L2. **(b)** Schematic summary of the six CAZy classes and their representative carbohydrate-active functions: Glycoside Hydrolases (GH), Glycosyl Transferases (GT), Polysaccharide Lyases (PL), Carbohydrate Esterases (CE), Auxiliary Activities (AA), and Carbohydrate-Binding Modules (CBM). **(c–e)** UMAP projections of the trained embedding spaces: L0 with CAZymes and non-CAZymes **(c)**, L1 with single-class CAZymes coloured by CAZy class **(d)**, and L2 with CAZymes coloured and labelled by CAZy family **(e)**.

Starting from the same pretrained ESM-2 foundation model^3^, we aligned the embedding space to three semantically defined resolutions using resolution-matched objectives (Fig. 2a). Membership-boundary and functional-grouping spaces were obtained by fine-tuning the ESM-2 backbone with binary and six-class classification objectives, respectively. The local-family space was obtained by applying a family-level Neighbourhood Components Analysis (NCA) metric-learning projection to ESM-2 representations. For compactness, we refer to these resolution-aligned spaces as Level 0, Level 1 and Level 2, respectively; implementation details are described in the Methods.

Each trained space emphasised a different level of biological organisation. Level 0 training produced a broad separation between CAZymes and non-CAZymes, consistent with a membership boundary around the target category (Fig. 2c). Level 1 training reorganised CAZymes into regions corresponding to the six major CAZy classes (Fig. 2d). The Level 1 geometry was uneven across classes, with large classes such as GT and GH occupying broad regions and smaller classes such as CE, PL and AA appearing as comparatively restricted regions. Level 2 training further fragmented the class-level into local family-level neighbourhoods, with numerous family-labelled subclusters visible within the broader CAZyme space (Fig. 2e). These resolution-matched spaces therefore provided the basis for asking which biological relationships become meaningful as neighbourhood relationships at each resolution.

### Membership-boundary embeddings selectively preserve target-category boundaries

We next asked whether the membership-boundary (Level 0) embedding space preserves the boundary between inside and outside the target category. As Level 0 supervision distinguishes CAZymes from non-CAZymes, we reasoned that sequences outside the CAZyme reference distribution should remain geometrically separated from the CAZyme manifold. We therefore projected three contrast-probe sets (denoting external proteins or perturbed sequences) into each trained CAZyme embedding space: shuffled CAZyme sequences, which preserve amino acid composition while disrupting sequence order; intrinsically disordered regions (IDRs), which are compositionally biased regions of proteins; and non-CAZyme proteins from those that are distinct to CAZy proteins (Fig. 3a). The shuffled and IDR sets serve as out-of-distribution (OOD) contrast probes: the shuffled set provides an artificial, composition-preserving sequence-order perturbation, whereas the IDR set provides a natural disordered-region contrast. The non-CAZyme set belongs to the supervised negative class for Level 0, although these sequences are not used during training. For the Level 1 and Level 2 spaces, this set serves as an external OOD probe. Separation from the CAZyme reference distribution was quantified using a per-sample silhouette score^19^, where higher values indicate probe separation from CAZymes and values near or lower than zero indicate significant mixing with the CAZyme sequences (Methods).

**Fig. 3.**
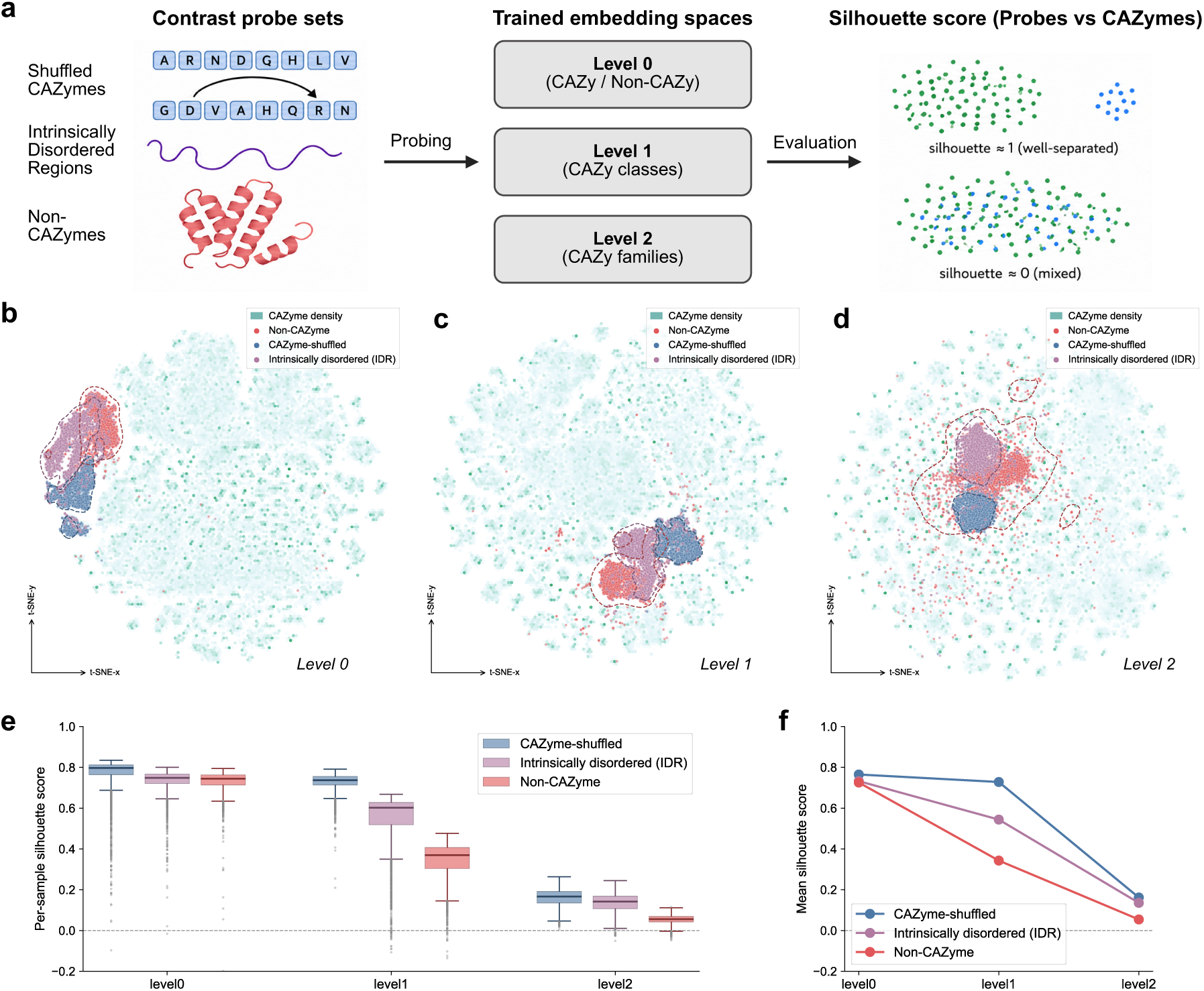
Contrast probe analysis across CAZyme embedding resolutions. **(a)** Schematic of the contrast probe analysis. Three probe sets were projected into the Level 0, Level 1 and Level 2 CAZyme embedding spaces: shuffled CAZyme sequences, generated by permuting amino acid order while preserving composition; intrinsically disordered regions (IDRs), representing compositionally biased sequences without stable globular structure; and non-CAZyme proteins from outside families. Separation from the CAZyme reference distribution was quantified using a per-sample silhouette score, where higher values indicate stronger separation from CAZymes and values near zero indicate mixing with the CAZyme manifold. **(b–d)** t-SNE projections of the contrast probes overlaid on the CAZyme reference density at L0 **(b)**, L1 **(c)** and L2 **(d)**. The CAZyme reference distribution is shown as teal density, and contrast probes are shown as coloured points with dashed outlines. **(e)** Per-sample silhouette score distributions for each probe type across L0, L1 and L2. **(f)** Mean silhouette score for each probe type across embedding resolutions.

At Level 0, all three probe sets remained separated from the CAZyme reference distribution, with the highest silhouette scores observed for shuffled CAZymes (Fig. 3b, e). This ordering was broadly maintained at finer resolutions, but the absolute degree of separation decreased as supervision shifted away from membership-boundary discrimination. Furthermore, the three probe types showed distinct patterns of boundary decay. Shuffled CAZymes remained relatively well separated from Level 0 to Level 1, indicating that class-level supervision still retains some ability to reject the most extreme sequence-order perturbations (Fig. 3c, e, f). IDRs showed a more gradual decrease in separation across Levels 0, 1 and 2. By contrast, non-CAZyme proteins showed the sharpest drop already at Level 1. This indicates that ordinary proteins outside the CAZyme category are no longer reliably separated once explicit CAZyme-versus-non-CAZyme supervision is removed. The magnitude of this effect may also reflect the breadth and heterogeneity of the CAZy functional-grouping resolution: CAZy classes span diverse carbohydrate-active functions, and some classes, such as Auxiliary Activities, contain particularly heterogeneous functional subtypes. Thus, class-level organisation can preserve the relationships among CAZymes, while providing a weaker boundary against other proteins outside the CAZyme category. At Level 2, silhouette scores decreased further across all probe categories, consistent with increased mixing between external probes and the CAZyme neighbourhood structure (Fig. 3d–f).

These results show that target category boundaries are selectively preserved in the membership-boundary space but progressively decay as the embedding objective shifts toward functional-grouping and local-family organisation. The strong Level 0 separation therefore reflects the necessity of explicitly learning the CAZyme-versus-non-CAZyme boundary, rather than an automatic property of all CAZyme-trained embedding spaces. Thus, proximity within a finer-resolution embedding space is insufficient on its own to imply membership in the target enzyme category, highlighting the need for a membership-boundary representation where category membership is the biological question of interest.

### Multi-domain proteins acquire resolution-dependent neighbourhood meaning

We next examined how embedding spaces represent multi-domain CAZymes associated with more than one functional class. Many CAZymes are modular proteins containing multiple catalytic or binding domains, and some are therefore annotated to more than one CAZy class. Such proteins provide a stringent test of neighbourhood semantics because their interpretation depends on the relationships between parent CAZy classes rather than the assignment to a single class. We therefore held out multi-domain CAZymes from the training dataset and projected them into the Level 0, Level 1 and Level 2 embedding spaces (Fig. 4a). We denote dual-class annotations using a vertical bar; for example, CBM|GH refers to proteins annotated to both Carbohydrate-Binding Module (CBM) and Glycoside Hydrolase (GH) classes, but this schema does not indicate order. For each multi-domain protein, we quantified its position relative to the centroids of its two parent classes using the Barycentric Alignment Score (BAS; Methods). BAS is defined from the distance between a protein and the midpoint of its two parent-class centroids, scaled by the radius from this midpoint to the parent centroids. Positive BAS values indicate that a protein falls within this parent-centroid midpoint region, a BAS score of zero indicates the boundary of this midpoint region, while a negative BAS value indicates placement outside it. We use this score together with the embedding visualisations to assess whether multi-domain proteins occupy neighbourhoods consistent with compositional positioning between their parent functional groupings.

**Fig. 4.**
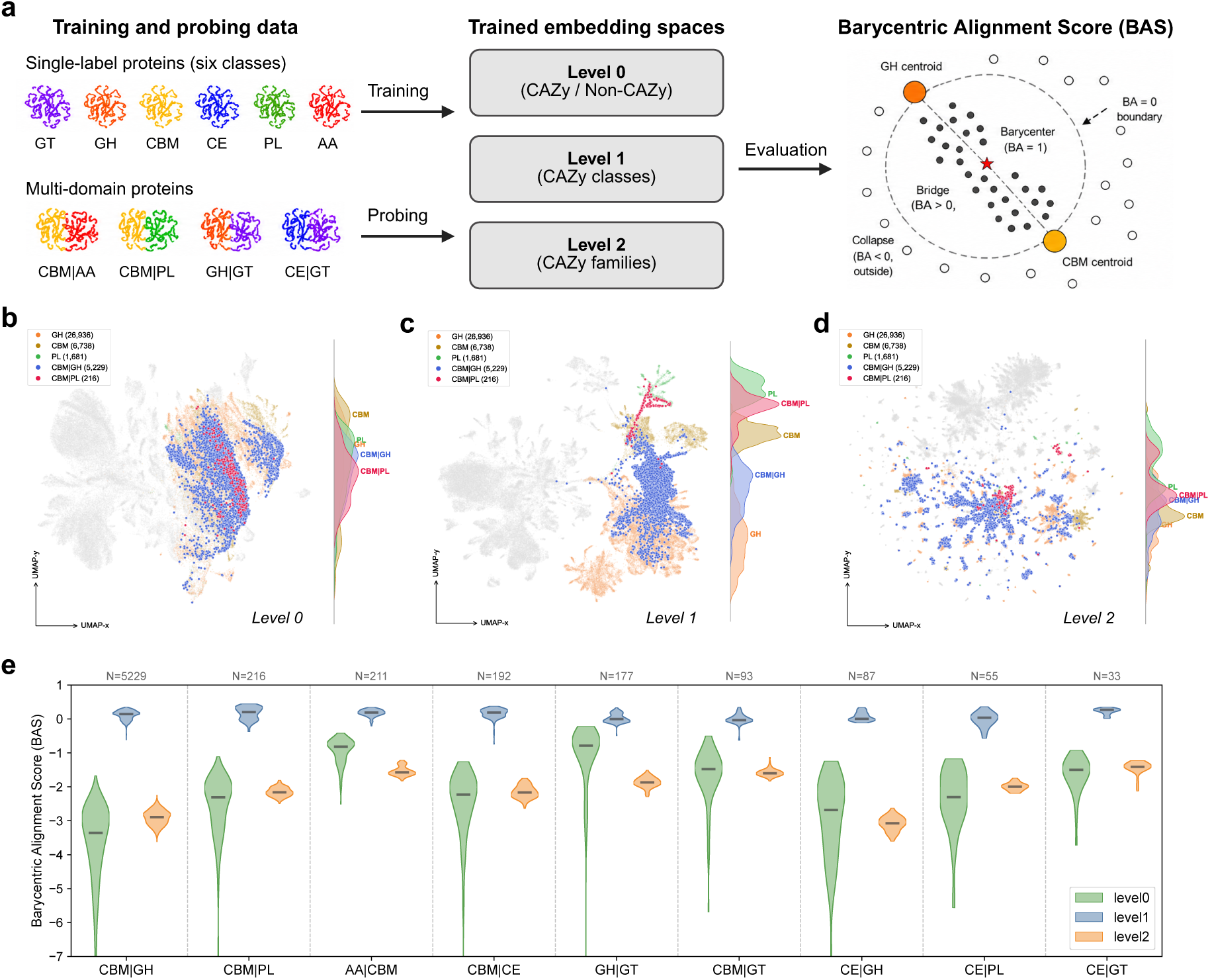
Multi-domain CAZymes across resolution-matched embedding spaces. **(a)** Schematic of the multi-domain CAZyme analysis. Multi-label CAZymes analysed here were defined as proteins annotated to two CAZy classes; these proteins were excluded from training and projected into the Level 0, Level 1 and Level 2 embedding spaces. The Barycentric Alignment Score (BAS) quantifies each protein’s position relative to the centroids of its two parent classes: BAS = 1 indicates placement at the parent-class barycentre; BAS > 0 indicates placement within the midpoint-centred region defined by the two parent centroids; and BAS < 0 indicates placement outside this region. **(b–d)** UMAP visualisations of two representative multi-domain class combinations, CBM|GH and CBM|PL, at L0 **(b)**, L1 **(c)** and L2 **(d)**, shown together with the corresponding single-class parent backgrounds. **(e)** Per-protein BAS distributions across nine multi-domain class combinations, ordered by sample size, in the L0, L1 and L2 embedding spaces.

At Level 0, multi-domain proteins remained inside the broad CAZyme manifold, but their parent classes were not sufficiently separated to define meaningful class-compositional geometry. For example, CBM|GH and CBM|PL proteins were embedded within the broad CAZyme region, where class-specific regions were poorly separated and intermediate positioning between parent classes could not be clearly interpreted (Fig. 4b). Consistent with this lack of parent-class organisation, BAS values were strongly negative and varied widely within each multi-domain group, such as CBM|GH or CBM|PL. This pattern indicates that membership-boundary supervision preserves CAZyme category membership, but does not provide a geometry that reliably constrains multi-domain proteins relative to their parent functional groupings.

In contrast, Level 1 embeddings resolved the major CAZy classes and placed multi-domain proteins along bridge-like regions between their corresponding parent-class neighbourhoods in the UMAP visualisation (Fig. 4c, e). BAS values shifted positively across class combinations and were centred around modestly positive values (Fig. 4e), supporting the visual observation that multi-domain proteins tend to occupy the parent-centroid midpoint region rather than falling outside it. This indicates that Level 1 supervision preserves compositional relationships between functional classes, with multi-domain proteins acquiring neighbourhood positions that reflect their dual functional annotation.

At Level 2, this class-compositional organisation was no longer maintained. Multi-domain proteins were redistributed into local-family level neighbourhoods rather than occupying positions between broad parent-class regions (Fig. 4d). BAS values became negative again, but with a pattern distinct from Level 0: within each multi-domain combination, the BAS score distributions were narrower, indicating a more consistently displaced position outside the parent-centroid midpoint region (Fig. 4e). This suggests that local-family level optimisation resolves multi-domain proteins according to local family-like structure, while no longer preserving the functional-grouping information that placed them between their parent classes at Level 1. Together, these results show that the same multi-domain proteins are interpreted differently across resolutions: as members of the CAZyme system at Level 0, as bridges between parent functional classes at Level 1, and as members of local family-level neighbourhoods at Level 2.

### Held-out families reveal the local semantics of family-level embeddings

We next asked whether proteins from families withheld from training can still form coherent local neighbourhoods in embedding space across the Level 0, Level 1 and Level 2. Under a conventional closed-set family classification system, proteins from sequence families absent during training cannot be assigned to their true family label. If Level 2 proximity reflects local-family level neighbourhood structure, rather than simply the embeddings modelling the families used for training, then held-out families should still form compact and distinct neighbourhoods at evaluation. To test this, we held out twenty-one CAZyme families spanning all six CAZy classes and projected them into the trained embedding spaces only during evaluation (Fig. 5a). For each held-out family, we computed a per-family Dunn index^20^, defined as the ratio between the nearest distance from the held-out family to any training-set family and the maximum pairwise distance within the held-out family (Methods). The Dunn Index matches the held-out family question: withheld families are expected to form discrete, compact neighbourhoods that remain distinguishable from their closest known-family context. High values indicate that a held-out family is both internally compact and geometrically separated from training families, whereas low values indicate merging with known families or internal dispersion.

**Fig. 5.**
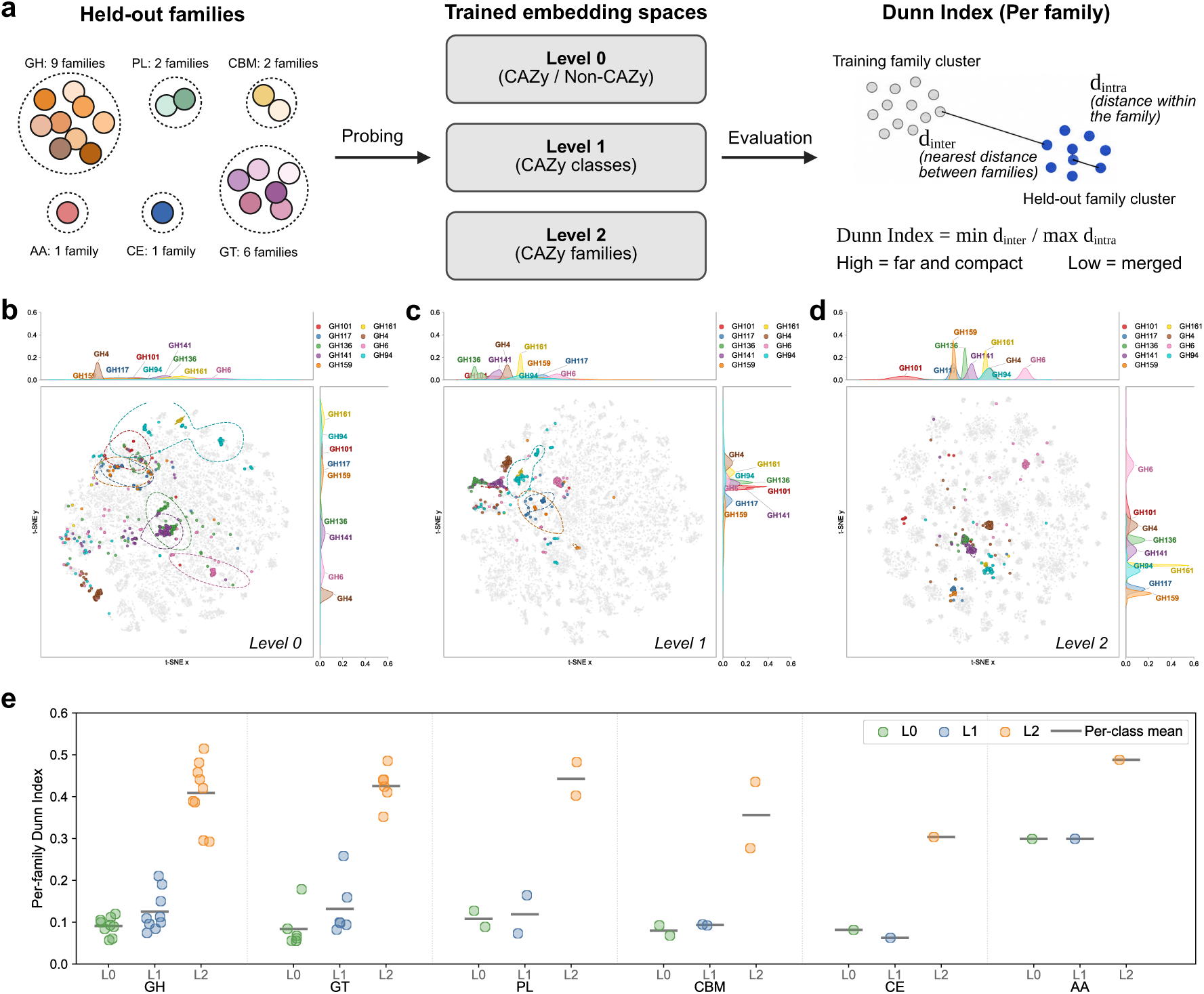
Held-out CAZyme family neighbourhoods across embedding resolutions. **(a)** Schematic of the held-out family evaluation. Twenty-one CAZyme families spanning six CAZy classes were excluded from training and projected into the Level 0, Level 1 and Level 2 embedding spaces at evaluation. The per-family Dunn index was computed as the ratio of the nearest distance from the held-out family to any training-set family to the maximum pairwise distance within the held-out family. **(b–d)** t-SNE projections of nine representative held-out GH families at L0 **(b)**, L1 **(c)** and L2 **(d)**, shown with training-set CAZymes as a grey background and marginal density distributions on each axis. **(e)** Per-family Dunn index values for held-out families across L0, L1 and L2, grouped by CAZy class; horizontal bars indicate per-class means.

At Level 0, held-out families were embedded among known CAZyme families rather than forming locally separated family neighbourhoods. Across all 21 held-out families spanning the six CAZy classes, Dunn index values were uniformly low at Level 0 (median, 0.089; range, 0.055-0.299), consistent with membership-boundary supervision preserving target category identity but not family-level organisation (Fig. 5e). Representative GH held-out families showed the same pattern clearly in the t-SNE visualisations (Fig. 5b). At Level 1, held-out families showed improved localisation within CAZy class regions compared with Level 0 (median, 0.099; range, 0.063-0.299; Fig. 5e). Representative families such as GH141 and GH161 began to display more compact structure at this resolution (Dunn index, 0.150 and 0.210, respectively; Fig. 5c). By contrast, Level 2 embeddings organised held-out families into compact, distinct neighbourhoods, with Dunn index values increasing sharply across all 21 held-out families (median, 0.424; range, 0.277-0.515; Fig. 5d,e). Thus, the median Dunn index increased approximately fivefold from Level 0 to Level 2, indicating a clear shift toward local-family neighbourhood structure. These results show that family-level proximity encodes local neighbourhood structure that generalises beyond the families used for training. Thus, sequences that would be treated as prediction errors under a closed family-label space can still occupy coherent neighbourhoods when examined at the appropriate embedding resolution. In combination with the reduced membership-boundary and functional-grouping signals observed at Level 2, this indicates that Level 2 embeddings are specialised for local sequence-family neighbourhoods, while broader biological relationships are represented less directly.

### Resolution-dependent neighbourhood semantics generalise across biological hierarchies and optimisation trajectories

Next, we investigated whether the neighbourhood semantics identified in CAZymes represent a system-specific phenomenon or a more general property of biological embedding organisation. To address this, we extended our analyses to peptidases, an independent hierarchical enzyme category organised across broad peptidase membership, catalytic type (aspartic, cysteine, glutamic, metallo, asparagine, mixed, serine and threonine peptidases and relate to the chemical groups responsible for peptide bond hydrolysis) and fine-grained peptidase families, with superfamily annotations, termed Clan in MEROPS, providing an additional intermediate reference level^17^ (Fig. 6a,b). Using the same resolution-matched strategy, we generated peptidase embedding spaces in which Level 0 represented membership-boundary resolution by distinguishing peptidases from non-peptidases, Level 1 represented functional-grouping resolution by organising peptidases by catalytic type, and Level 2 represented local-family resolution by organising peptidases by family (Methods).

**Fig. 6.**
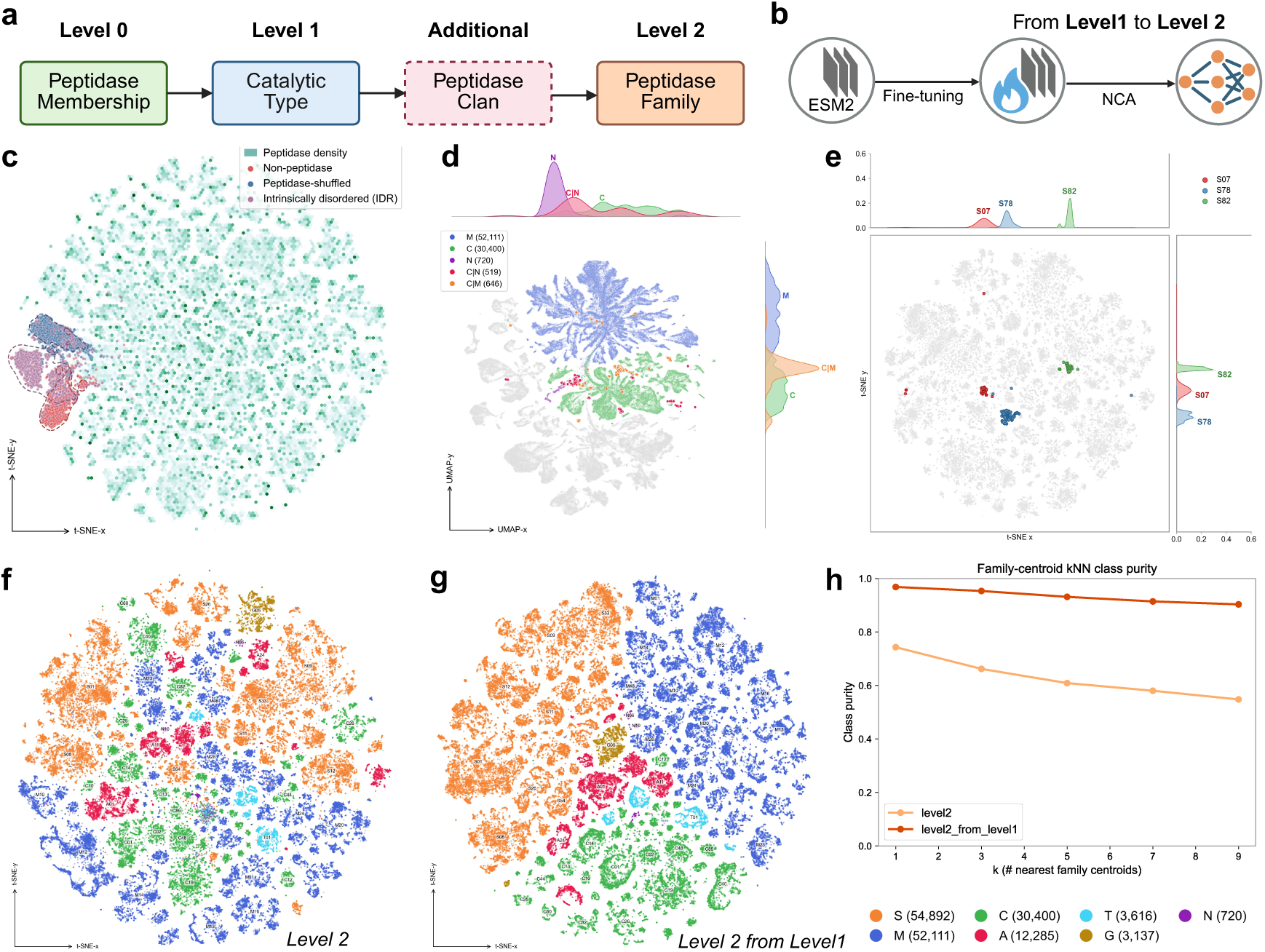
Peptidase embedding organisation across biological resolutions and training trajectories. **(a)** Peptidase annotation hierarchy, including peptidase membership (Level 0), catalytic type (Level 1, including pure types M, C and N and mixed types C|N and C|M), clan as an additional intermediate annotation, and peptidase family (Level 2). **(b)** Schematic of the sequential training trajectory from Level 1 to Level 2, in which ESM-2 representations are first fine-tuned using catalytic-type supervision and then aligned using Neighbourhood Components Analysis (NCA) at family level. **(c)** t-SNE projection of Level 0 peptidase embeddings with contrast probes, including shuffled peptidase sequences, non-peptidases and intrinsically disordered regions, overlaid on the peptidase reference density. **(d)** UMAP projection of the Level 1 embedding space showing pure catalytic types and mixed-type peptidases. **(e)** t-SNE projection of the Level 2 embedding space showing three representative serine-protease families, S07, S78 and S82, with marginal density distributions. **(f, g)** t-SNE projections of Level 2 peptidase embedding spaces obtained through two training trajectories: direct Level 2 family-level training from pretrained ESM-2 representations **(f)** and Level 2 family-level training after Level 1 fine-tuning **(g)**. Points are coloured by peptidase catalytic type. **(h)** Family-centroid k-nearest-neighbour catalytic-type purity across neighbourhood sizes for the two Level 2 training trajectories. Higher purity indicates stronger preservation of catalytic-type organisation among family-level neighbourhoods.

The peptidase system reproduced the resolution-dependent behaviours observed in CAZymes. At Level 0, contrast probes, including shuffled peptidase sequences, intrinsically disordered regions and non-peptidases, separated from the peptidase reference distribution, recapitulating the membership-boundary identity observed in CAZymes (Fig. 6c). Mean silhouette scores were highest among three levels for all three contrast probe types, including non-peptidases (0.6684), shuffled peptidase sequences (0.6110) and IDRs (0.5531). At Level 1, pure catalytic types formed distinct regions, while multi-domain peptidases were positioned between their corresponding parent catalytic types, mirroring the multi-domain bridge geometry observed for CAZymes (Fig. 6d). At Level 2, representative serine-peptidase families formed compact family-level neighbourhoods, consistent with local family structure emerging in an independent enzyme hierarchy (Fig. 6e).

Subsequently, we asked whether the path to a given resolution also shapes embedding geometry. We compared direct Level 2 training from pretrained ESM-2 representations with a sequential training trajectory in which the model was first aligned to Level 1 and then further optimised to Level 2 (Fig. 6f,g; Methods). Although both approaches aligned the embedding space to the same local-family resolution, they produced different family-level organisations. Direct Level 2 training recovered local family clusters but showed weaker preservation of catalytic-type structure, whereas Level 2 training after Level 1 alignment produced family clusters that remained more coherently organised by catalytic type. This difference was quantified by family-centroid k-nearest-neighbour catalytic-type purity (see Methods), which remained high across k for the Level 1-to-Level 2 trajectory but was substantially lower after direct Level 2 training (Fig. 6h), indicating that the same final resolution can preserve different higher-order relationships depending on optimisation trajectory.

## Discussion

Protein language model embeddings are often interpreted as maps of biological similarity, but our results suggest that their neighbourhoods should be interpreted according to the resolution at which the embeddings are organised. Across two representative yet independent enzyme hierarchies, the embeddings aligned to membership-boundary, functional-grouping and local-family resolutions emphasised distinct neighbourhood relationships. Thus, local proximity in an embedding space should not be assigned a fixed biological meaning; its interpretation depends on the biological relationship that the embedding geometry is organised to preserve.

This resolution-dependent paradigm also clarifies how embedding geometry relates to closed-label evaluation and optimisation history. Under a conventional closed-label setting, sequences from families absent during training cannot be assigned to their true family label, yet our analysis shows that they can still retain coherent neighbourhood structure at the appropriate embedding resolution. Conversely, coherent local structure should not be interpreted in isolation, because its biological meaning depends on its relationship to broader membership-boundary and functional-grouping organisation. In the peptidase trajectory analysis, embeddings aligned toward the same local-family objective retained different higher-order catalytic-type relationships depending on the optimisation path. Thus, embedding meaning is shaped by both the target resolution and the route through which that resolution is reached. Although this work focuses on protein language models, the concept of embedding resolution is not restricted to proteins. Any foundation model that maps biological entities into a continuous space faces a similar interpretability problem: neighbourhoods can reflect multiple possible scales of organisation. In DNA language models^21–23^, nearby sequences may be related through factors such as motif composition, gene content, mobile-element composition and/or genomic context. In single-cell and transcriptomic foundation models^24, 25^, neighbourhoods may reflect cell type, cell state, lineage, tissue context, perturbation response or technical covariates. The relevant biological resolutions differ across domains, but the interpretive question is the same: at what scale should local proximity be interpreted?

This also distinguishes embedding resolution from another active direction in protein representation analysis. Sparse autoencoder-based and related protein language model interpretability approaches have shown that latent features within models can correspond to biological concepts such as motifs, domains, binding sites or functional signals^26, 27^. More recently, representation-level approaches have begun to disentangle distinct biological constraints encoded within protein representations, including function- and stability-related constraints^28^. Embedding resolution addresses a different question: how the organisation of an embedding space shapes the biological interpretation of neighbourhood relationships. Under this view, the biological meaning of proximity is defined by the biological relationships that an embedding geometry is organised to preserve.

This framework also has implications for practical biological tools. Many embedding-based predictors are developed for a single task and evaluated mainly by benchmark performance^8, 29^. Our results suggest that representation design should be matched to the biological scale of intended use. Systems designed at different biological levels may require different embedding geometries, even when they are built from the same underlying foundation model. Moreover, the training trajectory itself can be used as a design choice: sequentially aligning embeddings through broader functional, structural or evolutionary groupings before local-family optimisation may help recapitulate hierarchical classification systems such as Pfam and CATH. In such settings, local neighbourhoods can support family- or domain-level interpretation while retaining information about higher-order organisation. In this sense, embedding resolution provides not only a framework for post hoc interpretation, but also a practical principle for designing task-specific and hierarchy-aware representations.

More broadly, our work opens several future directions that require further exploration. CAZymes and peptidases provide two independent enzyme hierarchies with well-developed annotations, but other functional protein groups may have weaker boundaries, less complete labels or different relationships between function, structure and evolution. Although we use ESM-2 as the foundation representation in this study, the resolution framework is decoupled from the specific foundation model. Applying the same analysis to other protein language models^4, 7^, structure-aware models^30^ or multimodal sequence–structure representations^9^ may reveal how different pretraining strategies shape resolution-dependent neighbourhood semantics.

Taken together, these findings reframe biological embeddings as resolution-dependent representations. Making resolution explicit shifts the question from whether an embedding space is broadly useful to which biological relationships its neighbourhoods are organised to preserve. This provides a basis for interpreting embedding neighbourhoods, designing task-specific representations and comparing biological foundation models across scales of interest.

## Methods

### CAZyme dataset collection and processing

We obtained 2,816,770 protein sequences from the CAZy database (2024 release)^16^. These were clustered at 50% sequence identity with MMseqs2^31^ and filtered for accession integrity, yielding 149,291 non-redundant sequences. Each sequence carries a hierarchical annotation: (i) a binary CAZyme designation, (ii) one of six catalytic or binding classes: glycosyltransferase (GT), glycoside hydrolase (GH), carbohydrate-binding module (CBM), carbohydrate esterase (CE), polysaccharide lyase (PL) and auxiliary activity (AA) — and (iii) the CAZy family (e.g., GH13, CBM4).

These sequences were processed in three successive steps. First, 9,086 sequences carrying only a catalytic or binding class label but lacking a family assignment were removed. Second, 6,431 sequences carrying annotations from two or more CAZy classes were set aside as a multi-domain probe set for alignment analysis (see Experiment 2). Third, 1,744 sequences from 21 held-out families were carved out as a held-out family probe (see Experiment 3). The remaining 132,030 single-class sequences were split in a class-stratified 7 : 1 : 2 ratio (seed = 42) into training (92,416), validation (13,205) and test (26,409) partitions.

As the negative set for binary (Level 0) fine-tuning, non-CAZyme proteins were drawn from Swiss-Prot^32^ (569,793 entries), clustered at 50% sequence identity with MMseqs2 (121,328 representatives) and decontaminated by removing sequences with detectable CAZyme homology (4,224 hits), yielding 117,104 non-CAZyme proteins split into training (81,972), validation (11,711) and test (23,421) partitions.

### Hierarchical resolution alignment framework

We used the 650-million-parameter ESM-2 protein language model^3^ as the basis for the resolution-alignment framework of three levels, each targeting a different tier of the hierarchical label system.

#### Level 0

The ESM-2 backbone was fine-tuned end-to-end for binary classification (CAZyme vs non-CAZyme) using a linear head over the pooled CLS-token representation, a cross-entropy objective, and a maximum sequence length of 1,022 residues (the remaining two positions are reserved for the CLS and EOS special tokens).

#### Level 1

The ESM-2 backbone was fine-tuned end-to-end for multi-class classification over the six CAZy classes, using a six-way linear head and a cross-entropy objective. Training followed the same protocol as Level 0 (sequence length, precision, optimizer), differing only in the classification target.

#### Level 2

A metric-learning projection network, a two-layer MLP (1,280 → 1,024 → 256) with L2-normalized output, was trained on the pooled CLS embeddings extracted from the pretrained (unfine-tuned) ESM-2 backbone using a single-label Neighbourhood Components Analysis (NCA) objective^33^ and an M-per-class batch sampler (M = 2).

### Embedding extraction and PCA dimensionality reduction

For every dataset partition we extracted pooled CLS-token embeddings (1,280-d) from the Level 0 and Level 1 fine-tuned backbones, and 256-d projected embeddings from the Level 2 metric-learning network.

### PCA dimensionality reduction

To produce a common, computationally tractable representation across all downstream analyses, each set of embeddings was passed through a two-stage pipeline: per-sequence L2 normalization followed by principal component analysis (PCA) retaining 100 components^34^. The pipeline was fit on the training partition and applied unchanged to all other partitions. The first 100 components recover 89.4%, 96.2% and 92.0% of cumulative variance for levels 0, 1 and 2 embeddings, respectively.

### Two-dimensional visualization

Projections of the 100-d PCA representation into two dimensions are computed with openTSNE^35^ for all per-class and contrast visualizations, and with UMAP^36^ for the joint training and multi-class visualization in Experiment 2. Both algorithms use PCA initialization for reproducibility.

### Experiment 1: contrast probes

#### Dataset construction

Three contrast sets were assembled to probe the geometry of the CAZyme embedding space. (i) The non-CAZyme test set (23,421 sequences, described above) was used directly; note that for the Level 0 embedding space, these sequences are in-distribution rather than out-of-distribution, since non-CAZyme proteins were part of the Level 0 training objective. (ii) A shuffled-sequence set (26,409 sequences) was generated by composition-preserving permutation of the CAZyme test set using esl-shuffle^37^, retaining per-sequence length and amino-acid composition while destroying primary-structure information. (iii) Intrinsically disordered proteins from DisProt^38^ (3,145 cluster representatives at 100% identity, length ≥ 21) served as an orthogonal sequence-composition control. For balanced comparison, each contrast set was subsampled to 3,145 sequences (seed = 42, matching the smallest set).

#### Visualization

In each level’s PCA representation, we computed a joint t-SNE projection of the CAZyme training set together with the three contrast sets. The training set is rendered as a hexagonal-bin density map, and each contrast set is overlaid as a scatter with a kernel-density contour enclosing the 90th density percentile.

#### Silhouette score

Geometric separation between each contrast set and the CAZyme training set was quantified with the silhouette score^19^, computed on the PCA representation. For each contrast sequence *i*, the silhouette is defined as:

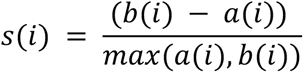

where *a*(*i*) is the mean Euclidean distance from *i* to all other sequences in the same contrast set, and *b*(*i*) is the mean distance from *i* to all sequences in the CAZyme training set. Values near +1 indicate clear geometric separation; values near 0 indicate overlap between the contrast set and the training distribution; values below 0 indicate that the sequence is, on average, closer to the CAZyme training set than to its own contrast group i.e., it is embedded within the training distribution rather than outside it. Per-sequence distributions and group means are reported for each level × contrast-set combination.

### Experiment 2: multi-domain probes

#### Dataset construction

From the 6,431 multi-class CAZyme sequences, we retained only those carrying exactly two CAZy class annotations and belonging to combinations with at least 30 members. Nine dual-class combinations passed this filter: CBM|GH (5,229), CBM|PL (216), AA|CBM (211), CBM|CE (192), GH|GT (177), CBM|GT (93), CE|GH (87), CE|PL (55) and CE|GT (33).

#### Visualization

A joint UMAP projection was computed on the PCA representation of the CAZyme training set together with the filtered multi-class set. Two representative combinations, CBM|GH and CBM|PL, were highlighted against a light-grey background, with marginal kernel-density panels comparing each combination to its parent classes. **Barycentric Alignment Score (BAS).** To quantify how a dual-class protein positions itself geometrically with respect to its two parent classes, we defined the per-sequence Barycentric Alignment Score (BAS). For a sequence *x* annotated with two classes:

- *c*₁, *c*₂ = the PCA centroids of the two parent classes, computed on the training set (single-class sequences only)
- *b* = (*c*₁ + *c*₂) / 2, the geometric midpoint of the two parent centroids
- *r* = (ǁ*c*₁ − *b*ǁ + ǁ*c*₂ − *b*ǁ) / 2, the mean centroid-to-midpoint radius

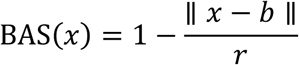

BAS = 1 places *x* exactly at the midpoint between its parent centroids; BAS = 0 places *x* at the mean centroid-to-midpoint distance; BAS < 0 places *x* outside the midpoint-centred region defined by its parent centroids. BAS is therefore a scale-normalized measure of how intermediate a dual-class protein lies between its declared parents in the embedding space. Per-combination BAS distributions are reported as kernel-density summaries with descriptive statistics.

### Experiment 3: held-out family probes

#### Dataset construction

To evaluate whether the resolution-aligned embeddings organise held-out families into geometrically coherent clusters, we constructed a held-out family probe set. For each of the six CAZy classes, families with between 20 and 200 training-set members were identified as candidates (mid-size: large enough for reliable evaluation, small enough to minimally perturb the training set). Approximately 10% of candidate families per class (at least one) were randomly selected (seed = 42) and their sequences removed from the training pipeline before splitting. This procedure yielded 21 held-out families totalling 1,744 sequences (AA: 1 family; CBM: 2; CE: 1; GH: 9; GT: 6; PL: 2). The main-text figure focuses on the GH class, with nine held-out GH families (GH4, GH6, GH94, GH101, GH117, GH136, GH141, GH159 and GH161) as visual exemplars.

#### Visualization

For each embedding level, a per-class t-SNE was computed on the GH-restricted PCA representation (training families plus the nine held-out families). All GH training sequences are rendered as a light-grey background, and each held-out family is displayed as a colored scatter with a 70th-density-percentile kernel-density contour. Per-axis kernel densities for each highlighted family are shown in the upper and right marginal panels.

#### Per-family Dunn index

Compactness and separation of each held-out family were quantified with a per-family Dunn index^20^. For held-out family *i*:

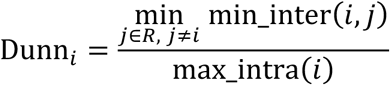

where *R* is the set of training families in the same CAZy class, min_inter(*i*, *j*) is the minimum Euclidean distance between any sequence of family *i* and any sequence of family *j*, and max_intra(*i*) is the maximum pairwise Euclidean distance within family *i*. High Dunn values indicate that family *i* is internally compact and well-separated from its nearest training-set neighbour. Distances are computed pair-by-pair to avoid materializing a full distance matrix.

### Peptidase replication

#### Dataset construction

To demonstrate that the framework generalizes beyond CAZymes, we assembled an independent peptidase dataset from the MEROPS database^17^. Starting from approximately 1.23 million entries in the MEROPS full-protein library, we retained only catalytically active classes: aspartic (A), cysteine (C), glutamic (G), metallo (M), asparagine (N), serine (S) and threonine (T), and removed inhibitors (I), unknown-mechanism entries (U), non-catalytic subunits (X), pseudogene annotations (1,417) and non-homologous fold entries (180,217), yielding 848,816 candidates. These were clustered with MMseqs2 at 50% identity (198,256 cluster representatives) and split per-family in a stratified 8 : 1 : 1 ratio into training (157,161), validation (19,552) and test (19,549) partitions.

Non-peptidase negatives were sourced from Swiss-Prot^32^ (121,328 entries after 50% clustering), decontaminated by removing sequences with detectable peptidase homology (MMseqs2 search, *e* < 10⁻⁵, identity ≥ 30%, coverage ≥ 50%), yielding 110,803 clean negatives split 8 : 1 : 1.

Two probe sets mirror the CAZy design: (i) a held-out family set of 1,994 sequences from 10 held-out MEROPS families (3 each from classes S, M and C; 1 from A; selected from families with 50–800 cluster representatives), and (ii) a cross-class two-catalytic-domain set of 1,525 proteins carrying domains from different MEROPS classes, analogous to the CAZy multi-class set. Additional contrast sets including non-peptidase sequences (11,081), shuffled peptidases (19,549) and the same DisProt set (3,145), were constructed following the same protocols as for CAZymes.

#### Pipeline replication

The entire three-stage fine-tuning pipeline (Level 0: binary; Level 1: seven-class; Level 2: NCA projection), embedding extraction, PCA reduction and Experiments 1–3 were executed independently on the peptidase data. All models were retrained from the pretrained ESM-2 base model.

### Metric-learning projection comparison

To isolate the contribution of supervised class fine-tuning to the quality of the downstream NCA projection, we compared two projection variants on the peptidase system.

**Level 2**: the NCA network (1,280 → 1,024 → 256) trained on pooled CLS embeddings from the pretrained ESM-2 backbone (no supervised fine-tuning).

**Level 2-from-Level 1**: the same NCA architecture trained on pooled CLS embeddings from the level-1-fine-tuned backbone.

Both projections share identical architecture, loss function, batch sampler and training hyperparameters; they differ only in whether the input features have been shaped by supervised class-level (Level 1) fine-tuning.

#### Visualization

Each projection was L2-normalized, reduced via PCA to 100 dimensions and visualized as a t-SNE coloured by MEROPS catalytic-type class.

#### kNN class purity

To quantify how well each projection preserves class structure at the family level, we computed MEROPS-family centroids in each 100-d PCA space (one centroid per family, averaging over all training sequences of that family; families with fewer than 5 members excluded). For each family centroid, we determined the fraction of its *k* nearest neighbouring centroids that share the same MEROPS class:

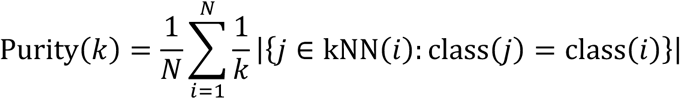

where *N* is the number of family centroids and kNN(*i*) denotes the *k* nearest centroids to centroid *i* (excluding itself), with *k* ∈ {1, 3, 5, 7, 9}. Higher purity indicates that families of the same catalytic type cluster together rather than interleaving with families of other types, reflecting a more class-coherent metric structure.

